# Transcriptional and synaptic regulation of NMDA glutamate receptor-mediated hippocampal plasticity and memory

**DOI:** 10.1101/2025.01.19.633774

**Authors:** Arnaldo Parra-Damas, Anna del Ser-Badia, Xavier Fernandez-Olaya, Ángel Deprada-Fernandez, Lilian Enríquez-Barreto, José Prius-Mengual, Judit Català-Solsona, Verónica Brito, José Rodríguez-Alvarez, Silvia Ginés, Antonio Rodríguez-Moreno, Carlos A. Saura

## Abstract

Synapse-to-nucleus signaling regulates activity-dependent synaptic plasticity underlying memory by linking N-methyl-D-aspartate (NMDA) glutamate receptors (GluN) to gene transcription mediated by the transcription factor cAMP-response element binding protein (CREB), but the underlying gene programs mediating potentiation at excitatory synapses are unknown. Here, we analyzed genome-wide chromatin immunoprecipitation sequencing (ChIP-seq) datasets of mouse and human CREB and the synaptonuclear factor CREB-regulated transcription coactivator1 (CRTC1) to identify relevant target genes and biological pathways coupling neuronal activity to synaptic function/plasticity. Our analyses indicate that CRTC1 specifically couples neuronal activity with synaptic plasticity by binding to conserved promoters of CREB target genes comprising inducible transcription factors (including *c fos*, *Crem*, *Npas4* and *Nr4a1*-*3*), and neuronal excitability and plasticity genes, including *Ntrk2*, *Homer1*, *Dlg4* (PSD-95) and the NMDA receptor subunit *Grin1* (GluN1). CRTC1/CREB target genes were highly enriched in gene ontology (GO) nuclear terms, including several members of the CREB family, and transcriptional modulators and repressors. Interestingly, GO enrichment and protein-protein interaction (PPI) network analyses revealed that genes mediating synapse-to-nucleus signaling (including most known synaptonuclear factors and direct interacting modulators) are collectively regulated by CREB/CRTC1, and that protein kinase C (PKC) is a key interactor of the CRTC1/14-3-3 complex at synapses. In agreement with these *in silico* analyses, we show that CRTC1 regulates synaptic activity-dependent phosphorylation and synaptic recruitment of GluN1 mediated by PKC in hippocampal neurons, and that PKC activation reverses NMDA receptor-mediated currents and long-term potentiation (LTP) deficits caused by CRTC1 silencing in the hippocampus. Consistent with genomics and functional data, morphological and behavioral analyses show crucial roles of CRTC1 on dendritic spine structure, plasticity, and hippocampal-dependent associative memory. Our results support a model in which neuronal activity and synaptic inputs are integrated in the nucleus through conserved CREB/CRTC1-regulated transcriptional programs sustaining global synapse-to-nucleus signaling pathways impacting on synaptic plasticity and memory.

## INTRODUCTION

One of the most remarkable properties of the brain is its capability to store, process and retrieve memories. Learning, memorizing and forgetting involve rapid and persistent enhancement or reduction of strength of excitatory glutamatergic synapses, processes known as long-term potentiation (LTP) and depression (LTD) (Lisman, 2017). These forms of neuronal plasticity are closely related to functional and structural modifications of synapses, especially at dendritic spines, the main sites of post-synaptic excitatory glutamatergic neurotransmission (Caroni et al., 2012). In the hippocampus, the prototypical LTP is mediated by glutamate N-methyl-D-aspartate (NMDA) receptors (GluN) and synaptic recruitment of a–amino-3-hydroxy-5-methyl-4-isoxazole propionic acid (AMPA) receptors (GluA), whereas synaptic LTD depends on glutamate-mediated removal of GluA from the post-synaptic compartments (Volianskis et al., 2015). Activity-dependent phosphorylation and synaptic recruitment of glutamate receptors induce and maintain synaptic plasticity in the CA1/CA3 hippocampus (Carroll and Zukin, 2002). Protein kinase C (PKC) mediates NMDA receptor gating and trafficking to synapses being required for NMDA-induced synaptic potentiation at CA3 synapses (Kwon and Castillo, 2008; Lan et al., 2001). Besides local synaptic mechanisms, long-lasting synaptic plasticity requires activity-dependent gene expression in the nucleus mediated by transcription factors and synapse-to-nucleus signaling mediated by synaptonuclear factors (Herbst and Martin, 2017; Kaushik et al., 2014). However, the global transcriptional programs by which synapse-to-nucleus signaling regulates glutamate-dependent synaptic plasticity are still unknown.

Memory depends on synapse-specific changes in neuronal excitability and synapse strength mediated, among others, by the transcription factor cAMP-response element binding protein (CREB) (Lisman et al., 2018). Surprisingly, the specific CREB-regulated transcriptional programs mediating neuronal excitability and synapse plasticity remain elusive. Synapse-to-nucleus communication is thought to mediate gene transcription underlying cognition and memory. Several synaptonuclear factors are transported from active synapses to the nucleus, allowing integration of synaptic inputs mediated by GluN NMDA receptors through activity-dependent transcription regulated by CREB (Kaushik et al., 2014; Parra-Damas and Saura, 2019). Accordingly, the synaptonuclear factor CREB-regulated transcription coactivator-1 (CRTC1) is critical for plasticity mechanisms underlying memory and emotional processes, including depression and addiction (Saura and Cardinaux, 2017). In neurons, concomitant calcium and cAMP signals induce CRTC1 dephosphorylation, nuclear translocation and transcriptional activity required for synaptic plasticity (Ch’ng et al., 2012; Kovács et al., 2007; Nonaka et al., 2014). In agreement with CREB-dependent synaptogenesis, dendritogenesis, synaptic plasticity and memory (Lee and Silva, 2009), CRTC1-dependent transcription mediates neuronal plasticity processes underlying memory (Nonaka et al., 2014; Parra-Damas et al., 2017a; Parra-Damas et al., 2014). Notably, deregulation of CREB/CRTC1-dependent transcription is associated with memory deficits in neurodegenerative disorders including Alzheimer’s, Parkinson’s and Huntington’s diseases [(Saura and Cardinaux, 2017), for a review]. Despite their critical role in synaptic plasticity and memory, how CREB/CRTC1-regulated transcriptional programs influence synapse-to-nucleus signaling to support structural and functional synaptic plasticity remained unexplored.

In this study, we used chromatin immunoprecipitation sequencing (ChIP-seq) to analyze the genome-wide occupancy profiles of CREB/CRTC1 in primary neurons, and explored its relevance in the regulation of synaptic mechanisms mediating plasticity and fear memory in the mouse hippocampus. We identified binding of CREB/CRTC1 to mouse and human ortholog target genes directly involved in synapse-to-nucleus communication, transcriptional regulation, synaptic plasticity, and neuronal excitability, including most of the known synaptonuclear factors, and the glutamate GluN1 (Grin1) receptor subunit -a key mediator of synapse-to-nucleus signaling. At the mechanistic level, we show that CRTC1 regulates synaptic activity-dependent PKC-mediated phosphorylation and synaptic recruitment of GluN1 in hippocampal neurons. Consistent with genomic data, functional, morphological and behavioral analyses show crucial roles of CRTC1 on dendritic spine structure and synaptic plasticity as well as fear-related memory, which are likely mediated through both distal transcriptional regulation and local synaptic signaling. Our results support a model in which neuronal activity and synaptic inputs are integrated in the nucleus through conserved CREB/CRTC1-regulated transcriptional programs sustaining global synapse-to-nucleus signaling pathways, impacting on synapse morphology, plasticity, and memory.

## RESULTS

### Conserved CREB/CRTC1 target genes mediate NMDAR synapse-to-nucleus signaling

To identify the genome-wide transcriptional programs regulated by CREB/CRTCs in neurons, we performed chromatin immunoprecipitation and next-generation sequencing (ChIP-seq) analyses of CREB, CRTC1, and CRTC2 in cultured primary neurons (11 DIV) in basal and neuronal activity conditions (**Fig. 1**). Induction of neuronal activity with forskolin (FSK) and potassium chloride (KCl), which promotes cAMP/Ca^2+^-mediated CRTC1 activation and nuclear translocation (Ch’ng et al., 2012; Parra-Damas et al., 2017b), potentiated genome-wide CRTC1 binding compared to vehicle control conditions (**Figs. 1A-G)**. Differential binding analysis (DBA) of the identified ChIP-seq peaks revealed significant binding of CREB and CRTC1, but not CRTC2 (**Fig. 1D-E and data not shown**). De-novo identification of transcription factor binding sites (TFBS) in the identified CREB and CRTC1 consensus peak sets (CPS) using JASPAR Enrich and FIMO/MEME, consistently revealed significant enrichment for CREB1 (**Fig. 1C**, data not shown). FSK/KCl treatment induced binding of CRTC1 to ∼2,000 sites distributed across ∼1,500 unique genes (**Data File S1; GEO accession number GSE131395).** We identified differential binding of CRTC1 between basal (vehicle) and stimulated (FSK/KCl) conditions (**Fig. 1E)**, which is reflected in the CRTC1 binding profile at proximal promoter regions, that is −3Kb to +3Kb from the transcription starting site (TSS) (**Fig. 1F**), whereas no differentially bound sites were identified in CREB ChIP-seq samples when comparing vehicle and FSK/KCl conditions (**Fig. 1A, data not shown**). However, the number of CREB peaks is much larger than the number of CRTC1 peaks and most CRTC1 peaks are also bound by CREB (**Data File S1**), while DBA between CREB and CRTC1 peaks reveal that most regions are preferentially bound by CREB (**Fig. 1D**). These results indicate constitutive binding of CREB and neuronal activity-dependent CRTC1 binding to target regions across the genome.

**Figure 1.**
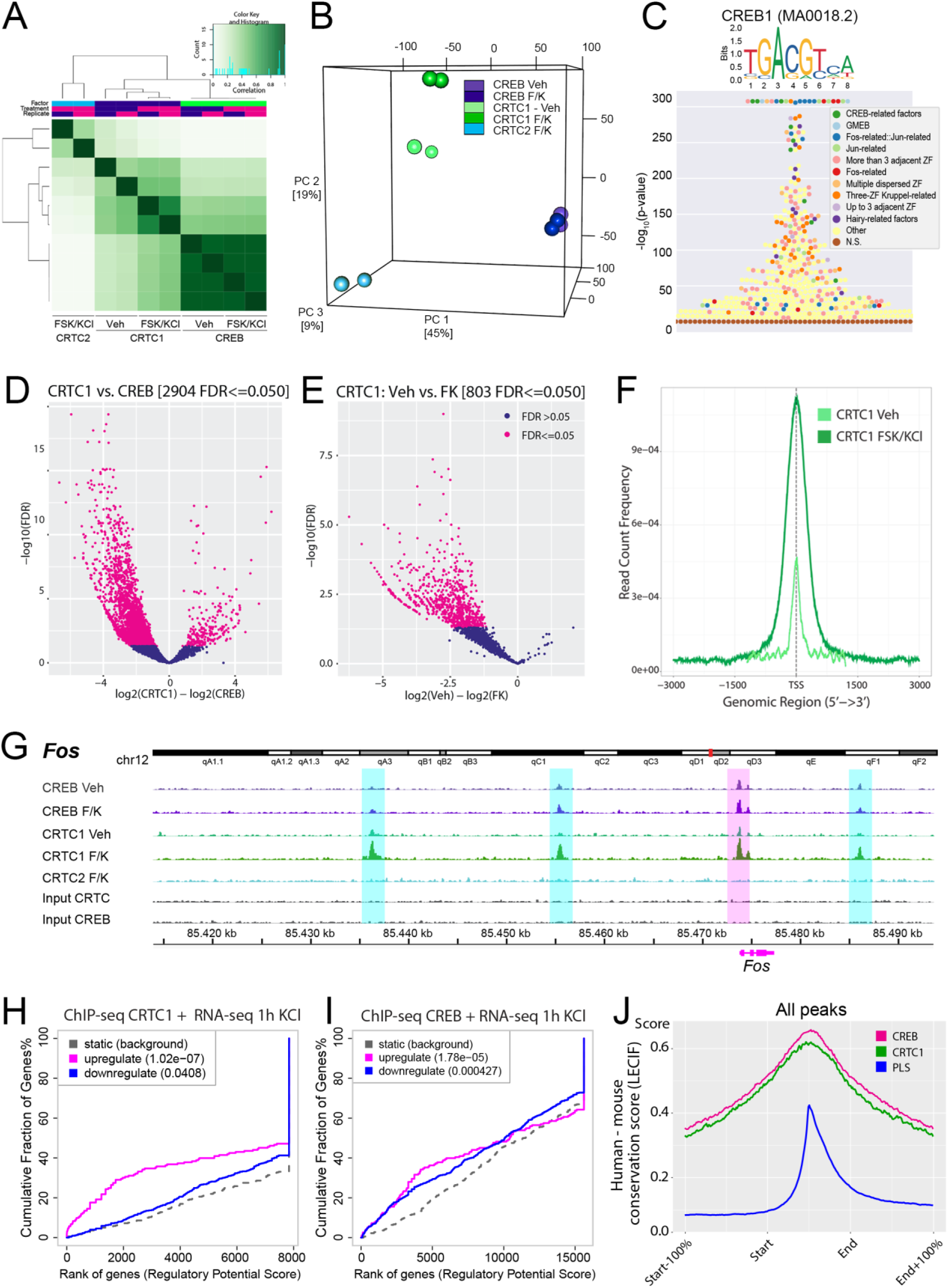
Activity-induced CRTC1 occupancy at CREB1 binding sites regulates neuronal transcription. **A-B,** Correlation heatmap (**A**) and 3D principal component analysis (PCA, **B**) of CRTC1, CRTC2 and CREB ChIP-Seq replicates, showing similar clustering of CREB but not CRTC1 samples in vehicle (Veh) and FSK/KCl (F/K)-treated conditions. **C,** JASPAR enrichment analysis of transcription factor binding motifs in the CREB/CRTC1 consensus peak set (CPS). **D**-**E,** Differential binding analysis (DBA) of the CRTC1 and CREB ChIP-seq experiments comparing CRTC1 and CREB ChIP-seq samples (**D**), as well as vehicle- and FSK/KCl-treatment conditions for CRTC1 (**E**). The volcano plot shows more differentially bound sites (red dots) for CREB compared to CRTC1 (negative log2(CRTC1)-log2(CREB) values in **D**), and only in CRTC1 ChIP from FSK/KCl (F/K)-stimulated neurons (negative log2(Veh)-log2(FK) values in **E**). **F,** Average genome-wide CRTC1 binding to proximal promoter regions (−3Kb to +3Kb from TSS) is induced upon neuronal activity. TSS: Transcription start site. **G,** Individual replicates of CREB, CRTC1, and CRTC2 ChIP-seq peaks identified around the *Fos* gene locus (bottom: in blue above the gene name), comprising 60 Kb upstream and 20 Kb downstream of the transcription start site. CRTC1 binding to the proximal promoter region (vertical magenta bar) was increased by FSK/KCl (F/K), while overall CREB binding was similar between basal (Veh) and FSK/KCl conditions. No specific binding of CRTC2 was detected in FSK/KCl-treated neurons. Known enhancers in the *Fos* locus are highlighted by blue bars. See **Data File S1**. **H-I,** Integration of RNA-seq with CRTC1 (**H**) and CREB (**I**) ChIP-seq data using BETA (Wang, et al., 2013. doi:10.1038/nprot.2013.150). RNA-seq data from KCl-depolarized mouse primary neurons was obtained from (Malik et al., 2014. doi:10.1038/nn.3808). **J,** Human–mouse conservation scores generated by neural network learning (LECIF; Kwon and Erst, 2021. doi:10.1038/s41467-021-22653-8) are higher in CRTC1 and CREB CPS compared to genome-wide mouse promoter like signature (PLS) sequences from ENCODE.

Binding of CREB and CRTC1 occurred at proximal and distal regions containing promoter-like signatures (PLS) and enhancer-like signatures (ELS) annotated in the ENCODE database (The ENCODE Project Consortium, 2020), and the corresponding target genes have consensus CRE sequences registered in the CREB target gene database (Zhang et al, 2005) (**Suppl. Figure 1A-B and Suppl. data file**). Integration of genomic conservation scores among vertebrates (PhyloP 60way) and between mouse and human genomes (LECIF) show high conservation scores in both proximal and distal regions (**Fig.1J**, **Suppl. Figure 1C** and **Suppl. data file**), indicating that the identified binding sites are conserved and correspond to functional regulatory regions.

Consistent with the role of CRTC1 as transcriptional co-activator of CREB, integration of published RNA-seq data obtained from mouse primary cortical neurons (DIV 7) treated with KCl (Malik et al., 2014) indicate that CRTC1 bound genes are preferentially up-regulated upon neuronal depolarization, whereas CREB-bound genes are both up-regulated and down-regulated (**Fig. 1H-I, Suppl.FigS1E-F**). Genomic conservation scores between mouse and human genomes (LECIF) indicate high conservation for the CREB and CRTC1 CPS, compared with mouse genome-wide promoter-like signatures (PLS) from ENCODE, suggesting that the identified binding sites are highly conserved and functional (**Fig. 1J**).

In agreement with the JASPAR enrichment identification of CREB1 consensus sequence in the CRTC1/CREB CPSs (**Fig. 1C**), enrichment analysis of TFs regulating the annotated CREB- and CRTC1-target genes using ChEA and ENCODE databases confirmed significant enrichment for CREB1 (**Suppl.Fig.S1D-E**). These results agree with recent CREB and CRTC1 ChIP-seq data from depolarized human iPSC-derived neurons, showing that CRTC1 regulates a subset of CREB-target regions (Boulting et al., 2021). Therefore, for subsequent analysis we generated a joint CREB/CRTC1 consensus peak set (CPS) containing 3620 peaks bound by either CREB or CRTC1 in at least 3 samples. As expected, and in agreement with the results obtained in the separate peak sets, enrichment analysis of TFs regulating the annotated genes in the joint CREB/CRTC1 CPS using ChEA and ENCODE databases, as well as identification of enriched TFBS in the CPS sequences, using JASPAR Enrich and FIMO/MEME, confirmed significant enrichment for CREB1 (**data not shown**).

To further confirm the functionality and conservation of the neuronal CREB/CRTC1-target regions we integrated our ChIP-seq data with independently published CRTC1/CREB ChIP-seq data obtained from depolarized human iPSC-derived neurons (Boulting et al., 2021), obtaining a significant overlap (revealed by hypergeometric and permutation tests) between our CREB/CRTC1 ChIP-seq peaks from mouse primary neurons and the CREB peaks from human iPSC-derived neurons after liftover of our mouse genomic sequences (mm10 to hg38), resulting in 2360 high-confidence binding sites conserved in human and mouse genomes (**Figure 2A** and **Suppl. data file**). Interestingly, the top 2000 peaks from each dataset exhibit ∼50% overlap (**Figure 2A, right**) suggesting a similar proportion of specific and common high-binding peaks between mouse primary neurons and human iPSC-derived neurons. Non-overlapping human and mouse peaks may reflect non/low-conserved binding sites or disparities in experimental procedures, including stimulation paradigms, but may also be due to distinct gene programs regulated in each cell type, since the human iPSC-derived neurons were characterized as GABAergic neurons, and our primary cortical mouse neurons are mainly developing excitatory neurons (data not shown). However, the profile of human-mouse LECIF conservation scores was higher in the 2360 overlapping peaks, compared to non-overlapping peaks or the total CREB peaks from human iPSC-derived neurons (**Fig. 2B**), suggesting that at least part of the differential binding between mouse and human regions may actually arise from lower sequence conservation.

**Figure 2.**
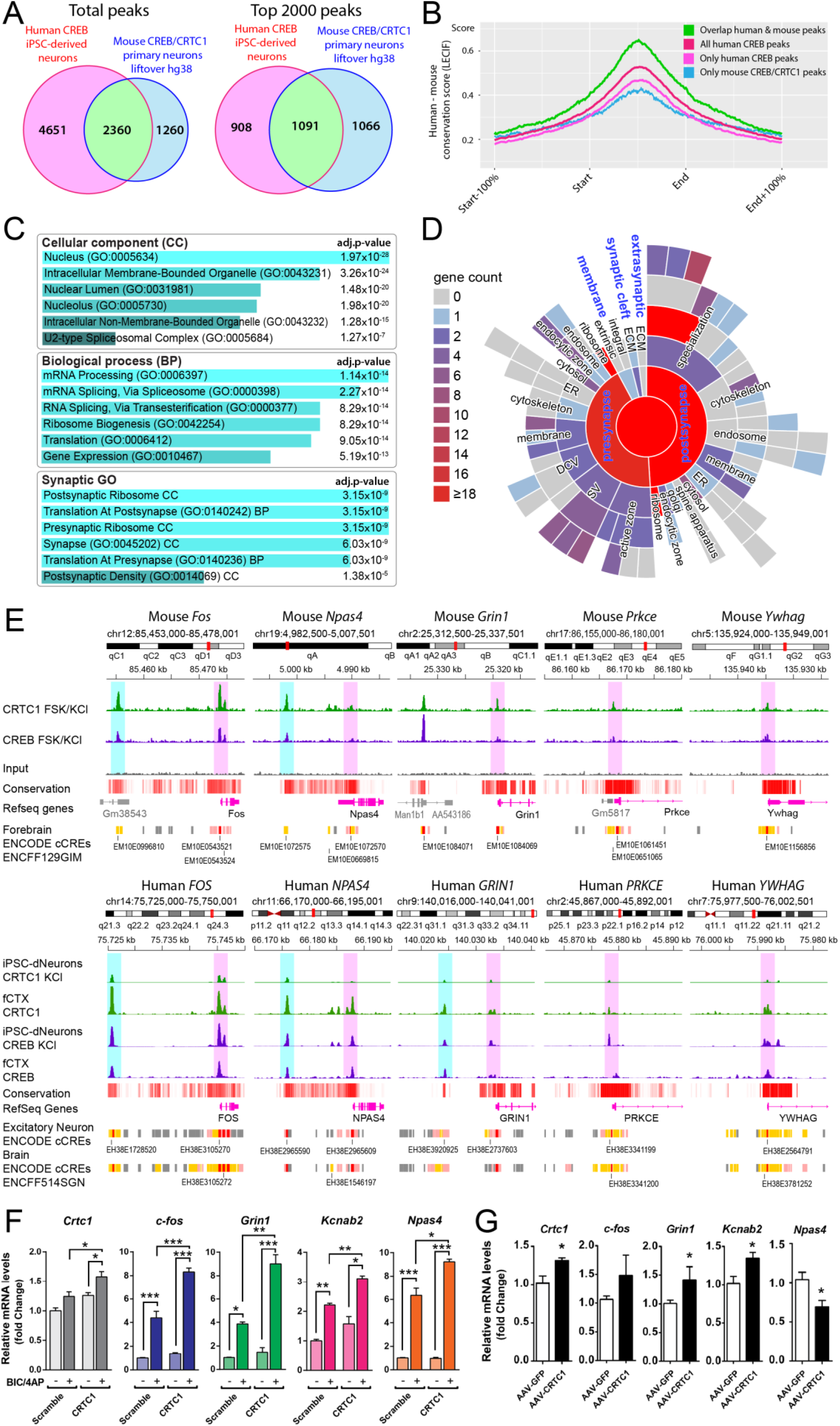
Conserved CRTC1/CREB target genes are enriched in nuclear and synaptic terms. **A,** Overlapping of mouse neuronal CREB/CRTC1 CPS (after LiftOver from mm10 to hg38) with published CREB ChIP-seq data obtained from human iPSC-derived neurons (Boulting et al., 2021. doi:10.1038/s41593-020-00786-1). **B**, Human-mouse LECIF conservation scores in the overlapping and non-overlapping total peaks obtained in **A,** (left). **C-D,** Gene ontology (GO) enrichment analyses of the conserved CREB/CRTC1 target annotated genes using Enrichr (**D**) (Xie et al., 2021. doi:10.1002/cpz1.90) and SynGO (E) (Koopmans et al., 2019. doi:10.1016/j.neuron.2019.05.002). **E,** Binding profiles of mouse (top) and human (bottom) CRTC1 and CREB ChIP-seq experiments showing proximal promoter regions (magenta bars) and active enhancers annotated in ENCODE (blue bars) of selected nuclear (*Fos*, *Npas4*) and synaptic (*Grin1*, *Prkce*, *Ywhag*) genes. Published human CREB and CRTC1 ChIP-seq data obtained from human iPSC-derived neurons and fetal cortical tissue (fCTX) was generated by (Boulting et al., 2021. doi:10.1038/s41593-020-00786-1). **F,** CRTC1 potentiates activity-induced gene expression in neurons. qRT-PCR analysis showing transcript levels of CRTC1 target genes in hippocampal neurons transduced with scramble- or CRTC1-lentivirus in the presence of TTX (-) or Bic/4AP (+). Values represent mean fold change ± SEM. *P < 0.05, ** P < 0.01, *** P < 0.0001, two-way ANOVA and Sidak’s multiple comparison test. **G,** CRTC1 affects gene expression in the adult hippocampus. qRT-PCR analysis of CRTC1 target genes in the hippocampus of 6-month-old mice transduced with adeno-associated virus (AAV)-GFP or-CRTC1. Values are mean fold change ± SEM. *P < 0.05, Mann-Whitney test.

We next focused on the 2360 conserved peaks-overlapped in human and mouse-, by annotating the corresponding human genes (hg38) and candidate cis-regulatory elements (cCREs) from ENCODE.

Interestingly, gene ontology (GO) enrichment analysis for cellular components (CC) and biological processes (BP) using Enrichr (Xie et al., 2021) revealed that conserved CREB/CRTC1-target genes are highly enriched in the nucleus (CC GO:0005634, adj.p-val = 1.974e-28) and in the BP terms Translation (BP, GO:0006412, adj.p-val = 9.025e-14), Gene Expression (BP GO:0010467, adj.p-val = 5.190e-13), and Cytoplasmic Translation (BP GO:0002181, adj.p-val = 8.228e-10), while enrichment analysis of synaptic GO terms annotated in SynGO (Koopmans et al., 2019) showed that 253 unique genes localize at synapses, exhibiting significant enrichment in the GO terms Synapse (CC GO:0045202, adj.p-val = 6.031e-9), Translation At Postsynapse (BP GO:0140242, adj.p-val = 3.153e-9), and Translation At Presynapse (BP GO:0140236, adj.p-val = 6.031e-9) (**Fig. 2C-D**). These results strongly indicate that CREB/CRTC1-regulated genes mediate both global neuronal gene expression and local translation at synapses. Accordingly, CREB/CRTC1 binding is detected at proximal and distal regions of genes encoding for nuclear transcriptional regulators and many synaptic genes (FOS, GRIN1, NPAS4, DLG4 [PSD-95], HOMER1, STX4/5, SNAP25, NRXN2, PRKCE, PRKCZ), including distal and proximal enhancer-like signatures (ELS) from brain and excitatory neurons annotated in ENCODE (**Fig. 2E**). In hippocampal neurons, synaptic activity induced with the GABAA receptor antagonist bicuculline (Bic) and the K+ channel blocker 4-amino-pyridine (4-AP) significantly increased transcript levels of c-fos, Grin1, Kcnab2 and Npas4, an effect potentiated by CRTC1 overexpression (P < 0.05, two-way ANOVA; **Fig. 1E**). Conversely, these genes were enhanced (c-fos, Grin1, Kcnab2) or decreased (Npas4) in vivo in the hippocampus of mice overexpressing CRTC1 using adeno-associated viral vectors (AAV2/10; **Fig. 1F**).

Since conserved CREB/CRTC1-regulated genes were enriched in the nucleus and also in the Postsynaptic Density (CC GO:0014069, adj.p-val = 0.1387 e-5), we next asked whether some of these genes are involved in synapse to nucleus signaling. Remarkably, we identified binding of CREB/CRTC1 to mouse and human ortholog target genes directly involved in synapse-to-nucleus communication, transcriptional regulation, synaptic plasticity, and neuronal excitability, including most of the known synaptonuclear factors (StNfs), and the glutamate GluN1 (*Grin1*) receptor subunit - a key mediator of synapse-to-nucleus signaling (**Supplementary** Fig.2). This observation prompted us to construct a gene regulatory network comprising all known genes encoding for StNfs and the CREB-target gene products regulating their expression (**Supplementary** Fig.2A). Highly connected (hub) CREB-target transcription factors (TFs) include Egr1 (regulating 9 StNfs and being targeted by 10 CREB-target TFs) and Yy1 (regulating 4 StNfs and being targeted by 12 CREB-target TFs and 3 StNfs). Finally, to identify synaptic proteins involved in CRTC1 synapse-to-nucleus signaling we performed protein-protein interaction (PPI) network analysis of CRTC1 partners at synapses using STRING and IntAct databases. PPI network analysis identified 14-3-3e and 14-3-3g isoforms as the main CRTC1 partners at synapses. Interestingly, we detected CREB/CRTC1 binding in promoter regions containing CRE consensus sequences of different 14-3-3 isoforms, including 14-3-3e, 14-3-3g, 14-3-3t and 14-3-3z, all of which are transcriptionally induced in depolarized primary neurons (Malik et al., 2014; **Suppl. data file**). To identify relevant mediators of CRTC1 synaptonuclear signaling we built a hierarchical PPI around CRTC1 and these 14-3-3 isoforms, restricted to synaptic proteins, resulting in 6 nodes corresponding to PKCe (PRKCE), SMAG1 (SAMD4A), LRRK2, PAK5, CDK16 and ABL1 (**Supplementary figure 2C**). Western blotting and qPCR analyses of mouse primary cortical neurons upon modulation of neuronal activity (TTX and FSK/KCl) and CREB function (Creb1 shRNA, and VP16-CREB) confirmed that most of the identified target genes are regulated by neuronal activity and/or CREB function (**Supplementary** Figures 3-4). These results suggest that CREB/CRTC1 act as master regulators of a highly conserved gene program mediating synaptic function and synapse-to-nucleus signaling, and that PKC, Egr1 and Yy1 may be key mediators of this program downstream of CREB. Accordingly, Egr1 expression is required for long term memory and facilitates synaptic plasticity (Katche et al., 2012; Maddox et al., 2010; Penke at al., 2013), whereas Yy1 regulates stress-induced transcription in excitatory neurons of the PFC (Kwon et al., 2022). Given the major role of NMDAR signaling on synapse-to-nucleus communication, and the importance of PKC for synaptic function and plasticity, we then explored the potential mechanisms linking CRTC1 and NMDAR/PKC-dependent plasticity.

### CRTC1 facilitates hippocampal synaptic plasticity by regulating NMDAR-mediated neurotransmission and structural plasticity

Since our ChIP-seq experiments suggested that CRTC1 regulates activity-dependent expression of genes mediating synaptic function, including Grin1 (GluN1), we next performed a detailed characterization of CRTC1’s effect on associative memory and functional and structural synaptic plasticity, including, LTP, LTD, NMDA and AMPA currents, and spine density and morphology (Fig. 3). AAV2/10 containing GFP (AAV-GFP), Crtc1-myc (AAV-CRTC1), scramble (AAV-Scr) or Crtc1 short hairpin interfering RNAs (AAV-ShCRTC1) were injected billaterally in the hippocampus of 4.5 month-old C57BL/6 mice. After 1.5 months, AAV-GFP or AAV-CRTC1 and -ShCRTC1 transduced mainly hippocampal pyramidal neurons and, compared to endogenous CRTC1, increased (∼4 fold; *P* < 0.0001) or reduced (∼50%; *P* < 0.001) CRTC1 levels, as previously reported (Parra-Damas et al., 2017a) (**Fig. S3**). Interestingly, CRTC1-myc was detected in the postsynaptic density fraction of AAV-CRTC1-injected mouse hippocampus (**Fig. S3).** Injected mice were evaluated for contextual fear memory at 6 months of age, prior to electrophysiological recordings. Statistical analysis showed a freezing time effect (GFP/CRTC1: F (2,33) = 21, *P* < 0.0001; ShScr/ShCRTC1, F (2,60) = 20.4, *P* < 0.0001; two-way ANOVA) with no significant group differences in basal and immediate freezing (*Post hoc*: *P* > 0.05. At 24 h, freezing responses were enhanced or reduced by CRTC1 overexpression and silencing, respectively (GFP *vs* CRTC1: *P* < 0.05; ShScr *vs* ShCRTC1 *P* < 0.05; two-way ANOVA; **Fig. 3A**). Electrophysiological recordings revealed robust long-term potentiation (LTP) at the Schaffer collateral-CA1 synapses in brain slices of memory-tested mice injected with GFP at 60 min (E-LTP: 145 ± 6%) and 180 min (L-LTP; 148 ± 6%; **Fig. 3B**). The magnitudes of E-LTP (173 ± 7%) and L-LTP (182 ± 6%) were increased 1.5 months after CRTC1 overexpression, whereas ShCRTC1 impaired both E-LTP and L-LTP (113 ± 8% and 94 ± 8%, respectively; *P* < 0.05, Student’s t-test; **Fig. 3B**). Next, we examined the effects of CRTC1 in long-term depression (LTD; 1 Hz, 30 min). This protocol elicited similar depression in GFP- and ShCRTC1-injected mice (GFP: 60 ± 5%, ShCRTC1: 54 ± 11%), but LTD magnitude was significantly increased in CRTC1-injected mice (45 ± 6%, *P* < 0.05, Student’s t-test; **Fig. 3C**).

**Figure 3.**
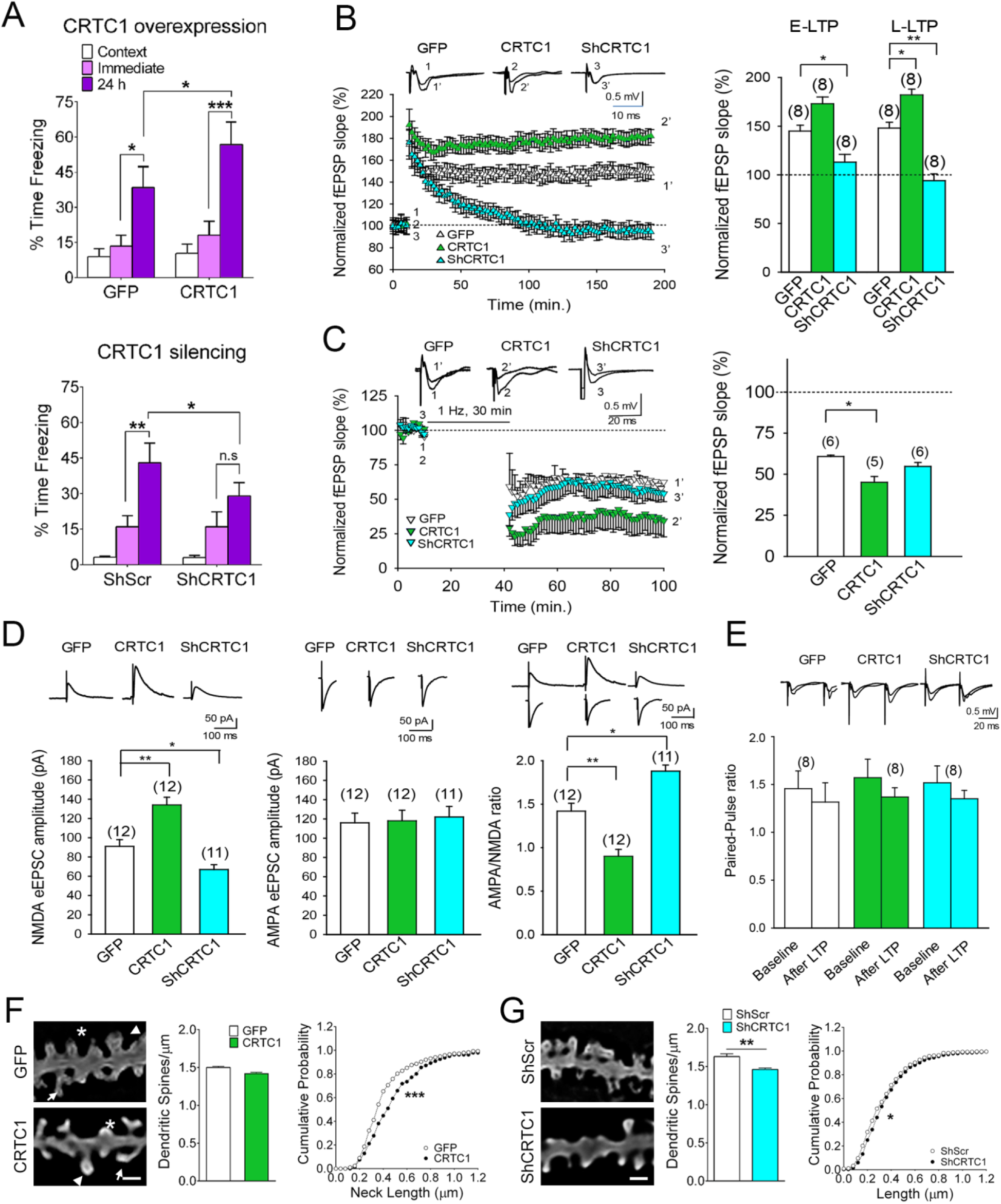
CRTC1 is critical for hippocampal NMDA receptor-dependent synaptic plasticity. **A,** CRTC1 is required for long-term associative memory. Mice intrahippocampaly injected with AAV-GFP or -CRTC1 overexpression vectors (n= 6-7/group; top) or AAV-ShScr or ShCRTC1 silencing (n=10-12/group; bottom) were tested in contextual fear conditioning. Values are % freezing ± SEM. \**P* < 0.05, ** *P* < 0.01, *** *P* < 0.0001, two-way ANOVA and Scheffé’s S *post hoc* tests. **B,** CRTC1 is critical for LTP at CA1 synapses. <u>Left</u>: Field excitatory postsynaptic potentials (fEPSP) slopes in 6-month-old mice injected with AAV-GFP (1, white symbols), -CRTC1 (2, green symbols) and -ShCRTC1-(3, blue symbols). Inset: traces show fEPSP before (1, 2, 3) and 180 min (1’, 2’, 3’) after TBS. <u>Right</u>: Average of E-LTP and L-LTP results of injected mice (n=4-5/group; slices in parentheses). Data represent mean ± SEM. \**P* < 0.05, ** *P* < 0.01, Student’s t-test. **C,** CRTC1 increases LTD in CA3-CA1 synapses. <u>Left</u>: fEPSP slopes before and after LTD induction (1 Hz, 30 min) in the CA1 hippocampus of 6-month-old mice expressing GFP (1, white symbols), CRTC1 (2, green symbols) and ShCRTC1 (3, blue symbols). Inset: traces show fEPSP before (1, 2, 3) and 60 min (1’, 2’, 3’) after LTD induction. <u>Right</u>: Normalized fEPSP slope results. Data represent mean ± SEM of several mice (n=4-5/group; slices in parentheses). \**P* < 0.05, Student’s t-test. **D,** CRTC1 enhances NMDA receptor-mediated excitatory postsynaptic currents. eEPSC amplitudes of NMDA (left; 10 μM NBQX and 20 μM bicuculline) and AMPA (middle; 10 μM D-AP5 and bicuculline) receptors and AMPA/NMDA ratio (right). Data represent mean ± SEM of multiple slices (parentheses) of injected mice (n=4-5/group). \**P* < 0.05, ** *P* < 0.01, Student’s t-test. **E,** Paired-pulse ratio does not change after LTP and is not affected by CRTC1. Inset: example traces during baseline and 180 after LTP induction. Data represent mean ± SEM. **F,** CRTC1 overexpression affects dendritic spine morphology but not density. <u>Left</u>: Deconvolved images of basal dendrites of Golgi stained-CA1 pyramidal neurons in 6-month-old mice injected 1.5 months earlier with AAV-GFP and -CRTC1. Images show stubby (*), small/thin (arrow) and large/mushroom (arrowhead) spines. Scale bars: 1 μm. <u>Middle</u>: Quantification of spine density in CA1 basal dendrites of mouse brains (n=3-5). <u>Right</u>: CRTC1 enlarges spine neck length. Global spine’s head areas and neck lengths in CA1 basal dendrites in mice injected with AAV-GFP (n=3) and AAV-CRTC1 (n=4). **G,** CRTC1 inactivation reduces the number of dendritic spines. <u>Left</u>: Images of dendritic spines of CA1 hippocampal neurons of 6-month-old mice expressing ShScramble (ShScr) or ShCRTC1 for 1.5 months. Scale bars: 1 μm. <u>Middle:</u> Quantification of spine density in CA1 basal dendrites (n=6 mice/group). <u>Right</u>: CRTC1 silencing increases spine neck length of CA1 neurons. n=6 mice/group.

Glutamate responses mediated by NMDA and AMPA receptors are essential for induction and/or maintenance of synaptic plasticity. We then investigated the role of CRTC1 on glutamate neurotransmission by using the whole-cell patch-clamp configuration. Interestingly, NMDA receptor-mediated current amplitudes were significantly elevated by CRTC1 (134 ± 8 pA) and decreased by ShCRTC1 (67 ± 5 pA) compared to GFP mice (90 ± 6 pA; *P* < 0.05, Student’s t-test), whereas AMPA receptor-mediated current amplitudes were similar in all groups (GFP: 116 ± 10 pA; CRTC1: 118 ± 11 pA; ShCRTC1: 122 ± 11 pA; **Fig. 3D**). Consistently, AMPA/NMDA ratios were significantly decreased and increased in CRTC1 and ShCRTC1 slices, respectively (*P* < 0.05, Student’s t-test; **Fig. 3D**). Paired-pulse facilitation ratios (PPR) before and after LTP induction revealed no significant changes among the groups indicating normal neurotransmitter release probability (*P* > 0.05), whereas input-output curves were slightly increased after CRTC1 inactivation (**Figs. 3E and data not shown;** *P* < 0.05). These findings demonstrate that CRTC1 modulates NMDA receptor-mediated currents in the adult hippocampus.

To investigate the role of CRTC1 in structural synaptic plasticity, we performed Golgi staining and dendritic spine morphology analysis in brains of 6-month-old AAV-GFP-or -CRTC1-injected mice (**Fig. 3F-G**). CRTC1 overexpression did not affect the number (GFP: 1.50 ± 0.01 spines/μm, CRTC1: 1.42 ± 0.02 spines/μm; *P* > 0.05, two-tailed Student’s t-test) or head areas of dendritic spines (*P* > 0.05), but it significantly enlarged spine necks of CA1 pyramidal neurons (*P* < 0.0001, Kolmogorov-Smirnov; **Fig. 3F**). Next, we analyzed the effects of CRTC1 genetic inactivation on basal dendritic spines of CA1 hippocampal neurons by using diolistic (DiI) labeling. This allowed quantification of synapse parameters in DiI neurons (red) that are mostly transduced with AAV-ShRNAs (*GFP*+; **Fig. S4A**). CRTC1 silencing caused a significant loss of spines (ShScr: 1.63 ± 0.04 spines/μm, ShCRTC1: 1.46 ± 0.02 spines/μm; *P* < 0.001, two-tailed Student’s t-test), and it increased neck length of large spines (*P* < 0.05, Kolmogorov-Smirnov; **Fig. 3G**). These results indicate that CRTC1 is critical for the maintenance and/or remodeling of dendritic spines in the hippocampus.

### Extranuclear CRTC1 regulates NMDA receptor phosphorylation and synaptic localization

Our findings of activity-dependent CRTC1 binding and regulation of *Grin1* promoter prompted us to examine the effects of overexpressing or silencing CRTC1 on glutamate receptors subunits levels.

Biochemical analyses showed that CRTC1 overexpression in the hippocampus did not affect total GluN1 levels, but it significantly increased phosphorylated GluN1 at Ser890 and Ser897 mediated by PKC and PKA, respectively (*P* < 0.05, one-way ANOVA; **Fig. 4A**), and the GluN2B/GluN2A ratio (*P* < 0.01, one-way ANOVA; **Fig. 4A**). By contrast, total and phosphorylated (Ser890) GluN1 levels were significantly decreased in the hippocampus of ShCRTC1-injected mice (*P* < 0.05, Student’s t-test; **Fig. 4A**). Importantly, AMPAR subunits (GluA1 and GluA2) and phosphorylated GluA1 (Ser831, Ser845), at two sites known to regulate GluA1 trafficking at the cell surface, were not affected by CRTC1 overexpression or inactivation (*P* > 0.05; **Fig. S5**). These results indicate that CRTC1 is required to maintain the levels and PKC-dependent phosphorylation of GluN1 (Ser890) in the adult hippocampus.

**Figure 4.**
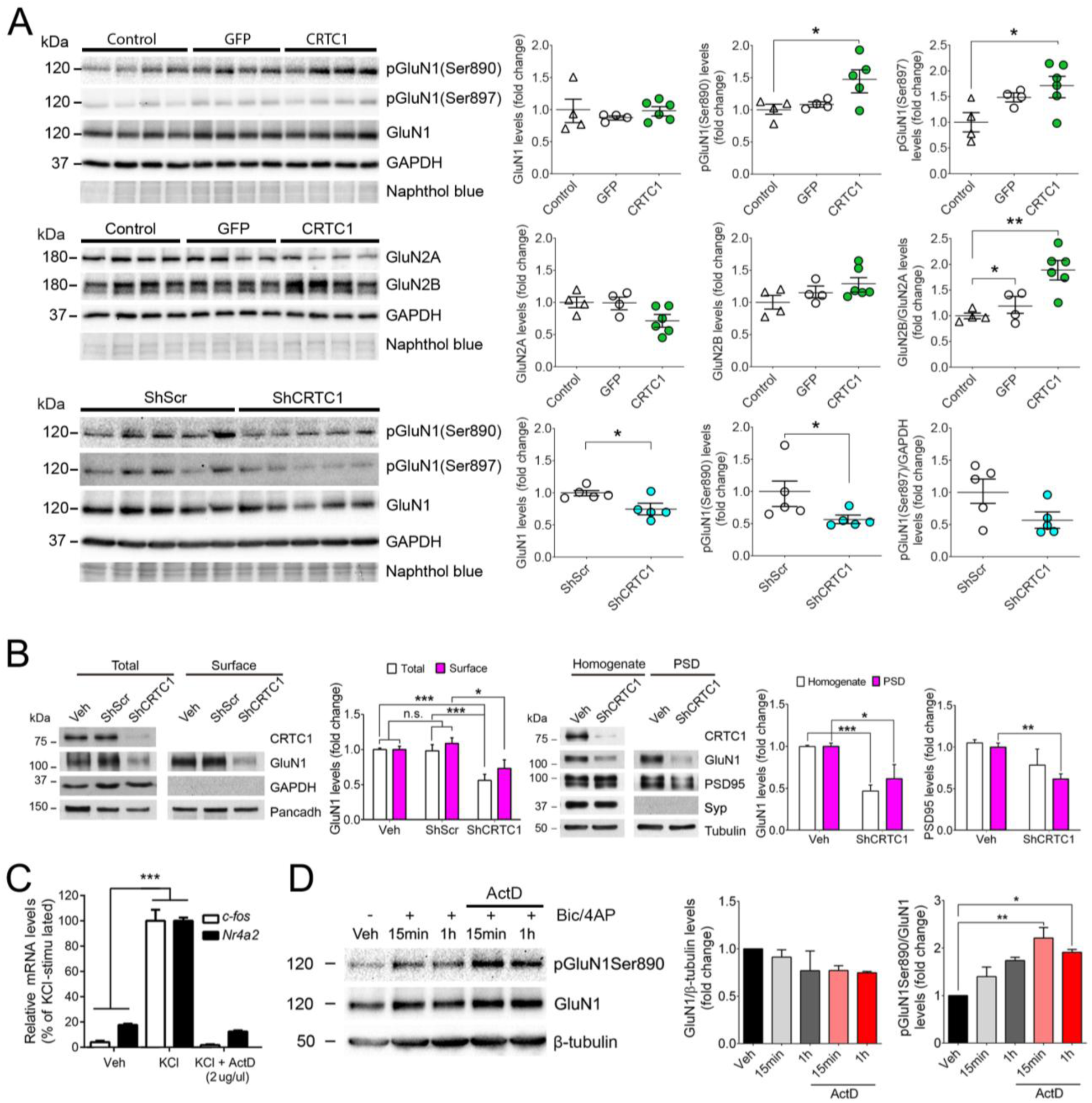
CRTC1 regulates GluN1 phosphorylation and synaptic localization in the hippocampus. **A,** CRTC1 regulates GluN1 phosphorylation in the hippocampus. Biochemical analysis of total and phosphorylated (p) GluN1 (Ser890, 897), GluN2A and GluN2B normalized to GAPDH in hippocampal lysates of 6 month-old control or AVV-GFP-, -CRTC1-, Scramble-or CRTC1 ShRNAs-injected mice. Data represent fold change ± SEM (n=4-5 samples/group). Statistical analysis was performed by one-way ANOVA followed by Bonferroni test or one tailed Student’s t-test. *\*P* < 0.05, *\*\*P* < 0.01 as indicated. **B,** CRTC1 is required for surface and synaptic localization of GluN1. <u>Left</u>: Biotinylation of surface GluN1 in transduced hippocampal neurons (18-20 DIV). <u>Right:</u> GluN1 and PSD95 in total homogenates and postsynaptic density fraction (PSD). Data represent protein levels ± SEM (n=4-10 cultures). Statistics was examined by two-way ANOVA followed by Tukey’s test. \**P* < 0.05, \*\**P* < 0.01, \*\*\**P* < 0.001. N.s: non-significant. **C,** Actinomycin D inhibits activity-dependent gene transcription. qRT-PCR analysis of *c-fos* and *Nr4a2* in hippocampal neurons treated as indicated. Data represent percentage RNA l ± SEM of n=4 independent cultures. \*\*\**P* < 0.0001 *vs* vehicle analyzed by two-way ANOVA. **D,** Actinomycin D does not prevent synaptic activity-induced GluN1 phosphorylation. GluN1 and pGluN1 Ser890/GluN1 levels in hippocampal neurons treated with vehicle (-) or Bic/4-AP +/- actinomycin D. Data represent mean ± SEM (n=3 experiments). \**P* < 0.05, \*\**P* < 0.01 as indicated analyzed by one-way ANOVA.

Biotinylation and subcellular fractionation assays in primary hippocampal neurons showed that CRTC1 silencing significantly reduced both total and surface GluN1, but not total PSD95, and it decreased GluN1 and PSD95 in purified postsynaptic density (PSD) fractions (*P* < 0.001-0.05, two-way ANOVA; **Fig. 4B**). No significant changes of these synaptic proteins were found between vehicle- and ShScramble-treated neurons (**Fig. 4B)**. To examine whether activity-dependent GluN1 phosphorylation and PSD95 dephosphorylation were dependent or independent of transcription, we performed treatments with actinomycin D (ActD), a chemical inhibitor of transcription. In our experimental conditions, ActD efficiently blocks *c-fos* and *Nr4a2* expression in KCl-depolarized neurons (*P* < 0.0001; **Fig. 4C)**. Interestingly, ActD did not prevent increase of phosphorylated GluN1 (Ser890)/GluN1 levels induced by synaptic activity for 15 min or 1 h (**Fig. 4D**; *P* < 0.05 vs vehicle). This result indicates that CRTC1/activity-dependent GluN1 phosphorylation mediated by PKC (Ser890) does not requires gene transcription, suggesting that CRTC1 could mediate local signaling mechanisms at synapses independent of its transcriptional regulatory role at the nucleus.

Activity-dependent CRTC1 nuclear translocation and transcriptional activity is mediated by Calcineurin-mediated dephosphorylation at several residues. We evaluated the effect of a phosphorylation-deficient CRTC1 mutant harboring three serine-to-alanine point mutations in the main phosphorylation sites mediating activity-dependent CRTC1 dephosphorylation and nuclear translocation (Ser64, Ser151 and Ser245), as well as two different constructs harboring point mutations in the nuclear localization signal (mNLS1 and mNLS2) (Ch’ng et al, 2015) (**Fig. 5A**). Surprisingly, the triple phosphorylation-deficient CRTC1 mutant exhibited impaired nuclear translocation and failed to induce CRE-transcriptional activity in the absence of endogenous CRTC1 (**Fig. 5C-E**). Conversely, both CRTC1 NLS mutants exhibited impaired nuclear translocation as expected, but surprisingly, were able to induce transcriptional activity when the endogenous CRTC1 was knocked down, to the same extent as the exogenous wild type CRTC1 construct (**Fig. 5C-E**).

**Figure 5.**
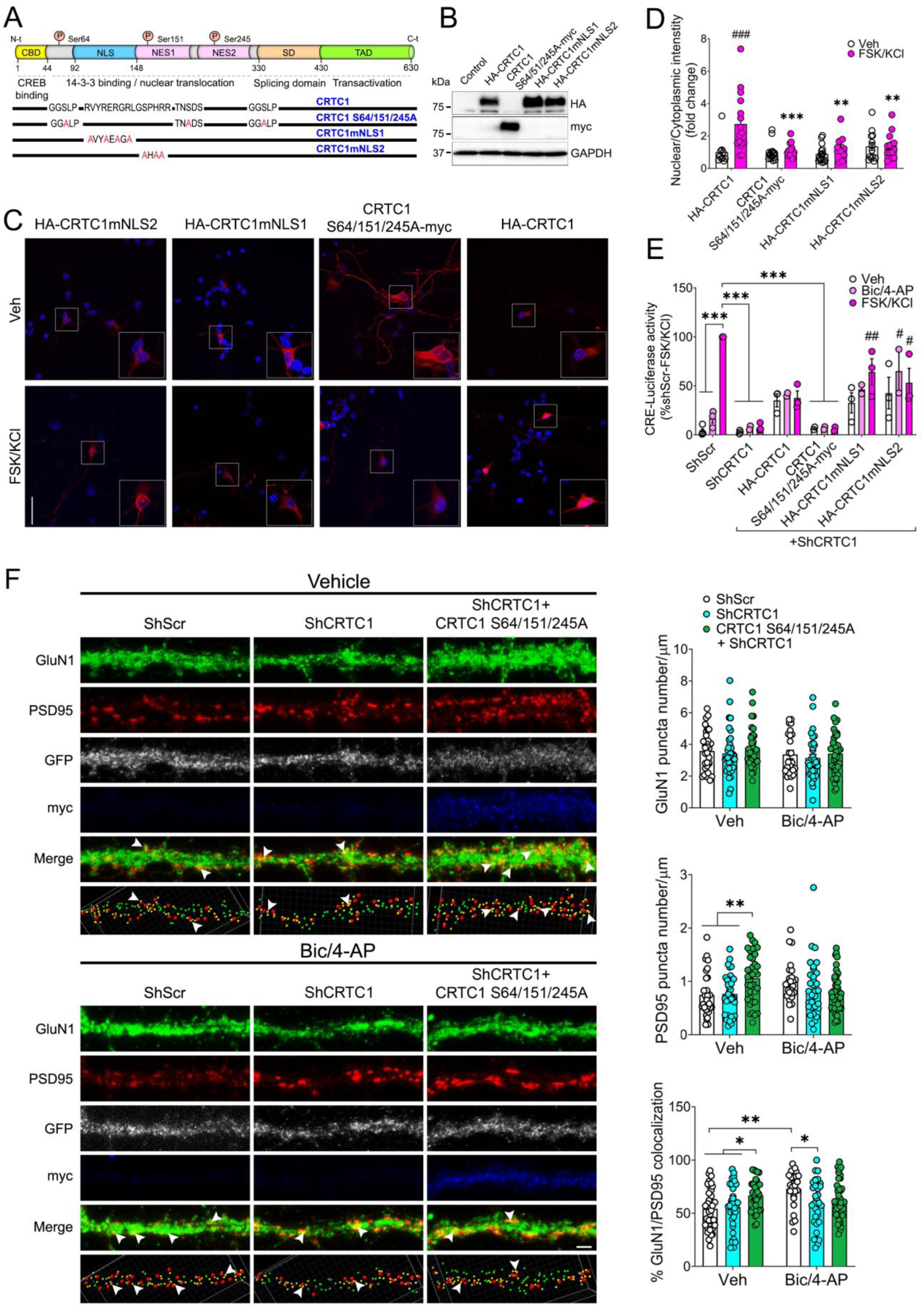
Extranuclear CRTC1 regulates activity-dependent GluN1 synaptic localization. **A,** Schematic representation of mouse CRTC1 (top) and CRTC1 S64/151/245A, CRTC1mNLS1 and CRTC1mNLS2 mutants. **B,** Western blot images of HEK293T cells transfected with wild-type and mutant CRTC1 constructs. **C,** Confocal images of transfected hippocampal neurons (7 DIV) with HA-CRTC1, CRTC1 S64/151/245A-myc, HA-CRTC1mNLS1 or HA-CRTC1mNLS2 (in red) treated with vehicle or FSK/KCl (20 µM, 30 mM) for 15 min. Scale bar: 40 µm. **D,** Quantification of nuclear/cytoplasmic CRTC1 intensity (n = 10-16 neurons from two independent cultures). Two-way ANOVA followed by Tukey’s post hoc test was used. \*\**P* < 0.01 and \*\*\**P* < 0.001 compared to HA-CRTC1 FSK/KCl; ^###^*P* < 0.001 compared to HA-CRTC1 Veh. **E,** CREB transcriptional activity analysis in cortical neurons (8 DIV) transduced with LV-ShScr or ShCRTC1 and transfected with the wild-type or mutant CRTC1. Neurons were treated with vehicle, Bic/4-AP (50 µM, 2.5 mM) or FSK/KCl (20 µM, 30 mM) for 4 h before analyzing the CRE-promoter luciferase activity. Data represent mean ± SEM of three independent experiments. Two-way ANOVA followed by Tukey’s post hoc test was used as a statistical test. \*\*\**P* < 0.001; ^#^*P* < 0.05 and ^##^*P* < 0.01 compared with ShCRTC1 FSK/KCl. **F, (Right)** Representative images of hippocampal primary dendrites (20 DIV) transduced with LV-ShScr or LV-ShCRTC1 alone or with LV-CRTC1S64/151/245A, pretreated with TTX (1 µM; 16 h) and stimulated with Bic/4-AP (50 µM, 2.5 mM; 1 h). Neurons were imaged for GluN1 (green), PSD95 (red), GFP (grey) and myc-tag (blue). Total GluN1 (green spots) and PSD95 (arrowheads/red spots) puncta per µm, and GluN1/PSD95 colocalization (arrowheads/yellow spots) were quantified. **(Left)** Bar graphs of the quantified spots. Data represent mean ± SEM (n = 24-42 dendrites from three independent cultures). Statistical analysis was determined by two-way ANOVA followed by Tukey’s post hoc test. \**P* < 0.05, \*\**P* < 0.01. Scale bar: 5 µm.

Because the phosphorylation-deficient CRTC1 triple mutant failed to induce CRE-transcriptional activity despite exhibiting robust expression and somatodendritic localization (**Fig. 5C-E**), we asked whether this mutant could regulate local activity-dependent changes in GluN1 signaling mediating synaptic plasticity, independent of transcriptional activity. Indeed, we observed that expression of the CRTC1 triple mutant -while knocking down endogenous CRTC1-increases the number of PSD95 puncta and GluN1/PSD95 colocalization in basal (vehicle), but not in Bic/4-AP-stimulated neurons (**Fig. 5F**), suggesting that CRTC1 exert local synaptodendritic effects regulating GluN1 synaptic localization in an activity-dependent manner.

### PKC activation mimics and occludes activity/CRTC1 effects on NMDA receptor-mediated synaptic plasticity

PKC-dependent GluN1 phosphorylation regulates cell membrane trafficking and synaptic potentiation in the hippocampus through still unclear mechanisms (Kwon and Castillo, 2008). By contrast, JNK1/PKCε-dependent PSD95 phosphorylation (Ser295) promotes synaptic/membrane PSD95 localization and promotes synapse number and potentiation (Sen et al 2016; Kim et al. 2007). To investigate how PKC activity influences GluN1 phosphorylation and synaptic localization, we next performed biochemical and fluorescence imaging analysis in mature hippocampal neurons. Pharmacological PKC activation with the phorbol ester 12-myristate 13-acetate (PMA) significantly increased phosphorylation of GluN1 (Ser890), an effect blocked by the PKC inhibitor GF-109203X (*P* < 0.05, one-way ANOVA and Bonferroni’s test; **Fig. 6A**). Notably, synaptic activity (Bic/4-AP;15 min) increased pGluN1(Ser890)/GluN1 without affecting the total levels (**Fig. 6A**). Confocal imaging analyses revealed that PKC activation increased synaptic GluN1 and PSD95 puncta, resulting in elevated GluN1/PSD95 colocalization in secondary dendrites of hippocampal neurons (%; *P* < 0.05%; two-tailed t-test; **Fig. 4C**). These results indicate that PKC activity induces GluN1 phosphorylation and GluN1/PSD95 synaptic colocalization in hippocampal neurons.

**Figure 6.**
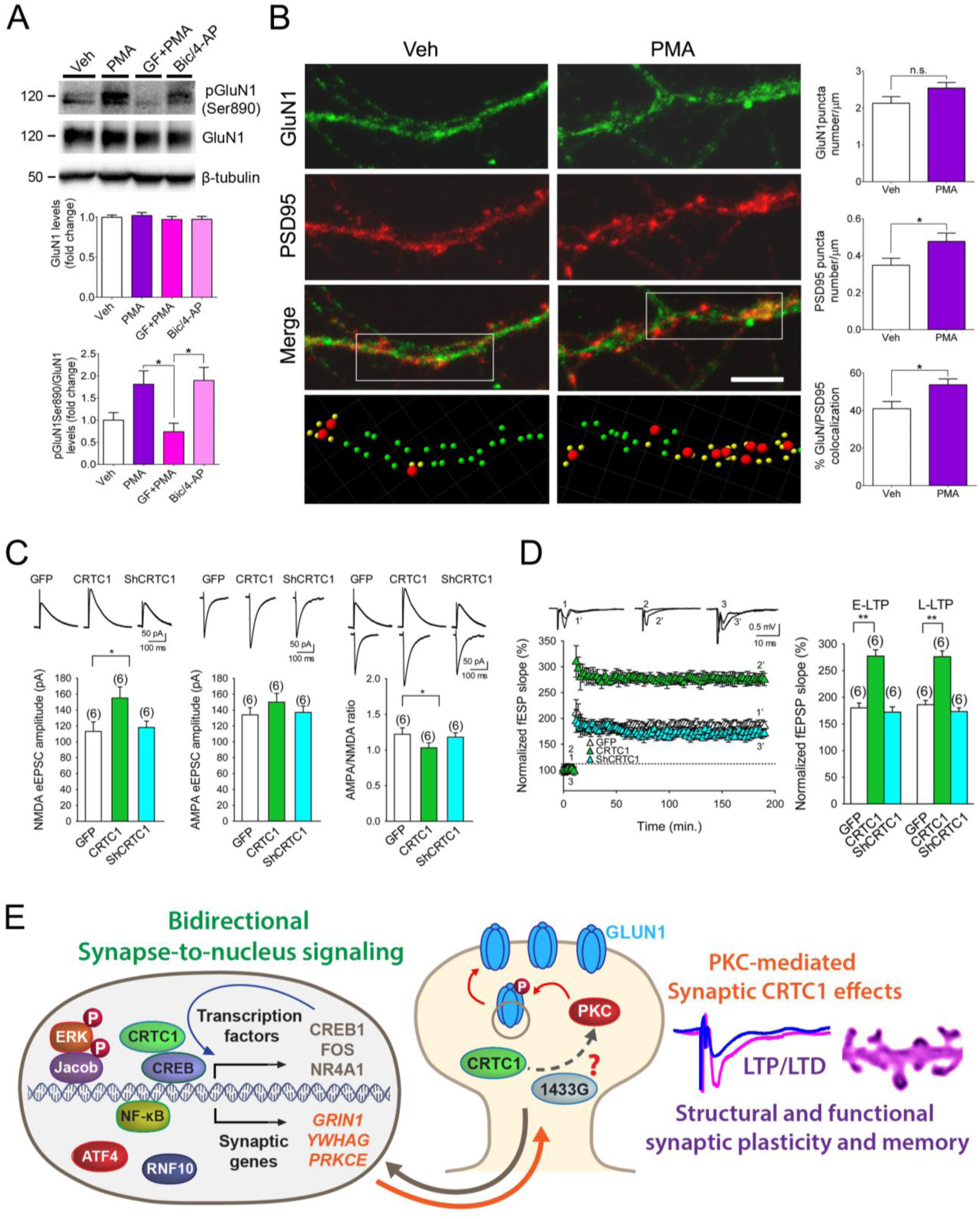
PKC activation mimics and occludes activity/CRTC1 effects on NMDA receptor-mediated synaptic plasticity. **A,** PKC enhances GluN1 phosphorylation. Total and phosphorylated GluN1 (Ser 890) in cultured hippocampal neurons treated with phorbol 12-myristate 13-acetate (PMA; 15 min) in the presence or absence of the PKC inhibitor GF-109203X hydrochloride (GF), or bicuculline (Bic) plus 4-amino-pyridine (4-AP). Data represent mean fold change ± SEM (n=6, independent cultures). Statistical analysis was determined by on-way ANOVA followed by Bonferroni’s test. \**P* < 0.05. **B,** PKC activation increases GluN1/PSD95 synaptic colocalization in hippocampal neurons. Confocal images of GluN1 (green), PSD95 (red) and GluN1/PSD95 (yellow) and quantification of merged spots in secondary dendrites of hippocampal neurons (20 DIV; n=19-23 dendrites/group; n=3 experiments) treated with vehicle or 12-myristate 13-acetate (PMA). Scale bar: 5 μm. Data represent mean ± SEM. Statistical analysis was determined by two-tailed t-test. \**P* < 0.05. N.s: non-significant. **C,** PKC activation reverses impaired NMDA receptor-mediated currents caused by CRTC1 silencing. eEPSC amplitudes of NMDA (left) and AMPA (middle) receptors and AMPA/NMDA ratio (right) in hippocampal slices from GFP-, CRTC1- and ShCRTC1-injected mice treated with phorbol 12,13-dibutyrate (PdBu; 1 μM). Representative current traces are showed at the top. Graphs represent mean eEPSC amplitude ± SEM of multiple slices (in parentheses; n=4-5 mice/group). \**P* < 0.05, Student’s t-test. **D,** PKC activation reverses hippocampal LTP deficits caused by CRTC1 silencing. <u>Left</u>: Field excitatory postsynaptic potentials (fEPSP) slopes monitored before and after a TBS plasticity protocol in brain slices of mice injected with GFP (1, white symbols), CRTC1 (2, grey symbols) or ShCRTC1 (3, black symbols) treated with PdBu. Representative traces (top) show fEPSP before (1, 2, 3) and 180 min after TBS (1’, 2’, 3’). <u>Right</u>: Average of E-LTP and L-LTP results. Graphs represent mean of normalized fEPSP slope ± SEM of multiple slices (in parentheses; n= 4-5 mice/group). ** *P* < 0.01, Student’s t-test. **E,** Schematic model of synaptonuclear-dependent synaptic plasticity. Local (synaptic) and distal (nuclear) mechanisms by which CRTC1 regulates synaptic potentiation. Abbreviations: NMDAR, N-methyl-D-aspartate receptor; LTP/LTD: long-term potentiation and depression; P, phosphorylation; PKC, protein kinase C.

Finally, we examined the role of PKC on CRTC1-dependent NMDA currents and plasticity in the hippocampus of AAV-GFP or AAV-CRTC1 and -ShCRTC1 transduced mice. Remarkably, pharmacological PKC activation with phorbol 12,13-dibutyrate (PdBu) increased NMDA current amplitudes in hippocampal slices of all groups, without affecting AMPA currents, and reversed the NMDA and NMDA/AMPA ratio amplitude deficits caused by CRTC1 silencing (GFP *vs* shCRTC1, *P* > 0.05; **Fig. 6A**). Furthermore, PdBu enhanced early and late LTP in all experimental groups (GFP, E-LTP: 180 ± 9%, L-LTP: 186 ± 8%; CRTC1, E-LTP: 277 ± 12%, L-LTP: 276 ± 11%; ShCRTC1, E-LTP: 172 ± 10%, L-LTP: 173 ± 7%; **Fig. 6D**, compared with **Fig. 3B**). Moreover, PdBu elevated significantly the magnitude of E- and L-LTP in CRTC1-overexpressing mice compared to GFP (*P* < 0.001) and ameliorated LTP impairments caused by CRTC1 inactivation (GFP *vs* shCRTC1, *P* > 0.05, Student’s t-test; **Fig. 6D**). These results demonstrate that PKC mediates CRTC1 effects on phosphorylation and synaptic recruitment of GluN1 upon synaptic activity, supporting NMDA receptor-mediated synaptic potentiation in the hippocampus.

## DISCUSSION

Synaptonuclear factors integrate and transduce neuronal signals received at distal synapses into transcriptional programs at the nucleus, but the synaptonuclear gene programs underlying potentiation at excitatory synapses are unknown. Here, we show that the synaptonuclear factor CRTC1 is as key mediator of neuronal excitability and synaptic plasticity by regulating CREB target genes including neurotransmitter receptors, synaptic proteins, voltage-gated ion channels and activity-induced transcription factors. Interestingly, CRTC1 binds to promoters of genes that act as negative regulators of CREB, such as *Crem*, *Atf4* and *Sik1,2* and *3,* suggesting a complex regulation of CREB target gene programs by activity-triggered negative feedback loops. It is likely that the overall genome-wide binding profile would not significantly differ between cortical and hippocampal neurons since CREB occupancy shares a high degree of peak overlap in these two neuronal types (Kim et al., 2010; Lesiak et al., 2013). Notably, neuronal activity induces CRTC1 binding to proximal promoter regions, and enhances transcription of *Grin1, Kcnab2* and *Npas4*. By contrast, CREB binding to the genome is similar in basal and neuronal activity conditions, consistent with our previous study showing constitutive CREB binding to specific promoter regions (Parra-Damas et al., 2017b). Our genome-wide ChIP-seq analyses identified CRTC1 binding to other activity-induced transcription factors (e.g., *Nr4a3*) not detected previously using conventional ChIP-qPCR assays (Parra-Damas et al., 2017b). Interestingly, a subset of CRTC1-regulated genes was not target of CREB, suggesting that some CRTC1-target genes may be regulated independently of CREB.

Besides the well-established function as transcriptional transductors of synaptic activity, synaptonuclear factors may potentially exert unknown local effects at synapses. Accordingly, CRTC1 facilitates synaptic plasticity by regulating expression, delivery and function of NMDA receptors at synapses. By contrast, inactivation of CRTC1 is detrimental for glutamate-dependent neurotransmission, synapse morphology and plasticity, and memory. These results suggest that this factor plays a critical role in both local and distal synaptic mechanisms mediating synaptic plasticity and memory, and its disruption leads to synaptic pathology and memory loss. Notably, CRTC1 acts as a fine-tuning regulator of structural and functional synaptic plasticity in the hippocampus. CRTC1 expression increases heads of stubby spines and neck length of small and large spines in CA1 hippocampal neurons, a result consistent with enlarged synapse heads induced by an active CRTC1 mutant (Nonaka et al., 2014). These synaptic changes are paralleled with enhanced synaptic efficacy, as evidenced by increased hippocampal NMDAR-dependent LTP and LTD. In this line, CREB also enhances NMDA receptor-mediated responses, excitability and LTP (Lopez de Armentia et al., 2007; Zhou et al., 2009). These results are also in agreement with previous findings indicating that CRTC1-dependent hippocampal LTP is mediated by transcriptional and epigenetic regulation of neuroplasticity genes (Kovács et al., 2007; Uchida et al., 2017). However, our results indicate that CRTC1 is required for maintenance, rather than induction, of early- and late-phases of LTP by regulating GluN1 phosphorylation and/or synaptic localization. Not only CRTC1 potentiates activity-dependent PKC-induced phosphorylation and synaptic delivery of GluN1 but also PKC activation efficiently reverses impaired LTP induced by CRTC1 inactivation.

It is plausible that CRTC1-mediated potentiation is due to enlargement of spine heads and/or necks, which elevates Ca^2+^ in spines and makes them preferential for induction of LTP (Noguchi et al., 2005). Indeed, longer and thinner necks permit larger Ca^2+^ levels in the head compartment, whereas larger spines, with shorter and wider necks, allow greater Ca^2+^ efflux into the dendritic shaft (Majewska et al., 2000; Noguchi et al., 2005). Spine size and synaptic strength are closely related as evidenced by LTP-induced spine enlargement and synaptic recruitment of AMPA receptors, a process that depends on NMDA receptors and CaMKII (Fortin et al., 2010; Matsuzaki et al., 2004). Interestingly, long-lasting potentiation requires the triheteromeric GluN1/GluN2A/GluN2B complex, whereas the transition from nascent (more plastic) to mature (stable) synapses is accompanied by a shift from GluN1/GluN2B to GluN1/GluN2A receptors (Gray et al., 2011; Volianskis et al., 2015). Remarkably, CRTC1-induced spine morphology changes are accompanied by a significant increase of GluN2B/GluN2A levels suggesting that CRTC1 changes dendritic spines towards a more dynamic and plastic phenotype. On the contrary, genetic CRTC1 inactivation causes loss and aberrant morphology of dendritic spines leading to synaptic potentiation impairments. The mechanisms underlying this synapse loss are unclear, but activity-induced synaptic weakening and spine shrinkage in CA1 neurons is mediated by NMDA and mGluR receptors (Oh et al., 2013). It is then possible that CRTC1-dependent NMDAR-mediated mechanisms are involved in both synapse maintenance and elimination.

The local synaptic mechanisms by which synaptonuclear factors regulate synaptic plasticity are still unclear. Our study shows that CRTC1 regulates synapse strength by regulating the expression, function and/or synaptic delivery of NMDA receptors. Synaptic activity induces CRTC1 binding to a proximal promoter region (<1 kb) of GluN1 increasing its expression *in vitro* and *in vivo*. In addition, CRTC1 potentiates activity-dependent PKC-induced phosphorylation of GluN1, whereas CRTC1 inactivation reduces total and synaptic GluN1 levels. The specific mechanisms by which CRTC1 mediates GluN1 phosphorylation are still unclear, but it may involve regulation of synaptic localization and/or activity of PKC or other related kinases. Nevertheless, activity-induced phosphorylation of GluN1 by PKC is independent of transcription. These findings are relevant because PKC facilitates the recruitment and mobility of NMDA receptors from extrasynaptic to synaptic sites (Fong et al., 2002; Groc et al., 2004; Tovar and Westbrook, 2002). Indeed, synaptic PKCα is activated rapidly by NMDAR and BDNF-TrkB signaling allowing integration of synaptic inputs into synaptic plasticity (Colgan et al., 2018). LTP also leads to rapid PKC-dependent phosphorylation and synaptic localization of GluN1, whereas PKC inhibition blocks NMDA receptor-mediated potentiation at CA3-CA1 synapses (Grosshans et al., 2002; Jeffrey et al., 2009; Kwon and Castillo, 2008). Considering that PKC-dependent GluN1 phosphorylation regulates its postsynaptic recruitment and synaptic potentiation [(Lan et al., 2001); this study], it is conceivable that CRTC1-dependent synaptic potentiation is mediated by GluN1 synaptic delivery and/or function. In support of this idea, PKC inhibition blocks activity-dependent GluN1 phosphorylation, whereas PKC activation enhances GluN1 phosphorylation and NMDA receptor-mediated currents and potentiation and reverses LTP impairments caused by CRTC1 deficiency. It is also plausible that CRTC1 regulates GluN1 synaptic trafficking through PSD95, whose synaptic delivery is decreased by activity in a phosphorylation-dependent manner (Kim et al., 2007). Our result indicating that synaptic PSD95 is reduced by CRTC1 silencing supports this idea. By contrast, CRTC1 does not affect GluA1 levels and AMPA neurotransmission. These results suggest that CRTC1 mediates synaptic potentiation in part by regulating activity-dependent expression, phosphorylation and/or synaptic delivery of GluN1.

Memory depends on changes in both neuronal excitability and synapse strength (Lisman et al., 2018). Our results indicate that CRTC1 regulates a subset of activity-dependent CREB target genes mediating synaptic plasticity and neuronal excitability, which are conserved in mice and humans. While identification of primate/human-specific neuronal gene programs are relevant for understanding distinct human cognition processes, we argue that focusing on gene programs that are conserved between human and biomedicine-relevant animal models is necessary to identify potential therapeutic targets.

CRTC1 may regulate neuronal excitability by controlling the expression, among others, of ion channels (e.g. *Kcnab2*) and inducible transcription factors such as *Npas4*, which regulates excitatory/inhibitory balance and synaptic plasticity during memory (Spiegel et al., 2014; Weng et al., 2018). The fact that CRTC1 regulates such a high number of downstream transcription factors (*c-fos*, *Nr4a1/2/3*, *Atf4*, *Crem*, *Npas4*…) indicate that CRTC1 acts as a first-line master regulator of activity-induced transcription in neurons. These results may be highly relevant for cognitive and neurodegenerative disorders in which CREB/CRTC1 deregulation is associated with memory deficits (Parra-Damas and Saura, 2019). Thus, altered *Grin1* expression is associated with several brain disorders, such as schizophrenia, and Alzheimer’s and Parkinson’s diseases (Hemby et al., 2002; Jacob et al., 2007). Of relevance, genetic variants in NMDA receptors and *PKC* correlate with hippocampal memory performance in humans (de Quervain and Papassotiropoulos, 2006), and mutations in the CREB coactivator *SRCAP* were recently identified in Alzheimer’s disease patients (Vardarajan et al., 2017). Whether these genetic variants affect transcription and synaptic plasticity by regulating synapse-to-nucleus signaling merits further investigation. Future studies are necessary to elucidate the local synaptic functions of these factors, so we can understand the role of synapse-to-nucleus signaling in brain physiology and pathology.

## ACKNOWLEDGMENTS

We are grateful to Jean-René Cardinaux (University of Lausanne, Swizerland) for providing the CRTC1-myc cDNA, Toh Hean Ch’ng (Nanyang Technological University, Singapore) for providing the CRTC1mNLS1 y CRTC1mNLS2 constructs, and Ángel Barco (Instituto de Neurociencias de Alicante, Spain), for providing the VP16-CREB and VP16-Control plasmids. We thank C. Gutiérrez, M. Castillo, N. Barba and S. Mendizuri for excellent technical assistance in cell culture, histology and imaging procedures, and members of our lab (O. Recasens, N. Mendizuri, C.M. Soto-Faguás and L. Ortega) for technical and experimental assistance. We thank the UAB Servei d’Estabulari, Viral Vector Production (UPV), Servei de Genòmica i Bioinformàtica (SGB) and Histology and Microscope INc facilities of UAB for technical support. This study was supported by research grants from Ministerio de Ciencia e Innovación (Spain) MCIN/AEI/10.13039/501100011033 with FEDER and “NextGenerationEU/PRTR” funds (SAF2016-80027-R, PID2019-106615RB-I00, PID2022-137668OB-I00 and PDC2021-121350-I00 to CAS; PID2019-107677GB-I00 and PID2022-136597NB-I00 to ARM, and PID2020-11751ORB-I00 and PID2023-151925OB-I00 to JRA), Instituto de Salud Carlos III (CIBERNED CB06/05/0042), Generalitat de Catalunya (2021 SGR00142), Junta de Andalucía (CVI-7290 to ARM), BrightFocus Foundation (A2014417S), and Alzheimer’s Association (AARF-24-1313365). A.P-D. has been supported by the following fellowships from Ministerio de Ciencia e Innovación: Juan de la Cierva-Incorporación (IJC2019-042468-I), and Ramón y Cajal (RYC2022-037843-I). AdS is supported by a predoctoral fellowship from Generalitat de Catalunya (2018FI_B00858). JCS is supported by Ayudas para la Formación de Profesorado Universitario fellowship, Ministerio de Educación y Cultura (FPU14/05392). A.D-F. and X.F-O. are supported by predoctoral FPU (FPU/02486) and FPI (PRE2020-092926) fellowships from Ministerio de Ciencia e Innovación.

## Author contributions

APD and CAS conceived and coordinated the study. APD, AdSB, LEB, JPM, JCS, VB, OU, JRA, SG, ARM and CAS designed and conducted experiments. APD, LEB, JRA, SG, ARM and CAS discussed the data and wrote the paper.

## Data and Materials availability

Materials are available on request. ChIP-seq data was deposited in the GEO database (GSE131395).

## MATERIALS AND METHODS

### Experimental Design

The objective of this study is to discern the CRTC1-dependent mechanisms mediating functional and structural synaptic plasticity and hippocampal-dependent memory. The study comprises the following experimental procedures: 1) Genomics analyses by using CRTC1, CRTC2 and CREB ChIP sequencing of the whole mouse genome in cultured neurons; 2) Validation and quantification of specific activity-dependent CRTC1 target genes in hippocampal neurons; 3) Role of CRTC1 in glutamate receptor-mediated hippocampal structural and functional (LTP and LTD) synaptic plasticity using AAV-mediated gain-of-function and loss-of-function (i.e. ShRNA) approaches; 4) Examination of CRTC1-mediated regulation of NMDA and AMPA glutamate receptors (expression, localization, phosphorylation…) in the mouse hippocampus and cultured hippocampal neurons; 5) Discerning the molecular mechanisms by which CRTC1 regulates synaptic delivery and phosphorylation of GluN1 in hippocampal neurons.

### Mice

All C57BL/6 mice were male and maintained in standard conditions of 12 h light/dark cycle with food and water available *ad libitum* at the Animal Core Facility of the Universitat Autònoma de Barcelona. Animal procedures were performed under institutional and national regulations approved by the Animal and Human Ethical Committee of the Universitat Autònoma de Barcelona (Protocols CEEAH 2896, DMAH 8787) in accordance with the experimental European Union regulations (2010/63/EU).

### Primary neuronal cultures

Cortical and hippocampal neurons were obtained from embryonic (E15.5) or postnatal (P0) mouse embryos (C57BL/6). Cortices and hippocampi of mouse brains were dissected and transferred to a sterile centrifuge tube containing 10 ml of Krebs buffer (120 mM NaCl, 4.8 mM KCl, 1.2 mM KH_2_PO_4_, 25 mM NaHCO_3_, 14.3 mM glucose, 0.3% bovine serum albumin and 0.03% Mg_2_SO_4_) and centrifuged (300 x *g*, 1 min). The pellet was resuspended in Krebs buffer containing 0.025% trypsin, incubated at 37 °C (10 min) and then stop solution (Krebs solution plus 0.052% trypsin inhibitor, 0.0008% DNAse and 0.03% MgSO_4_) was added. After centrifugation (300 x *g*, 1 min), the pellet was dissociated in Krebs solution containing 16% of stop solution and filtered through a nylon mesh. The filtered cell suspension was then transferred to a tube containing Krebs solution plus 0.03% MgSO_4_ and 0.0014% CaCl_2_ and centrifuged (250 x *g*, 5 min). The supernatant was discarded and the cell pellet was resuspended in B27/glutamine-supplemented Neurobasal medium. Neurons were counted in a hemocytometer using trypan blue. Neurons were seeded in poly-D-lysine-coated 24 well-dishes (30,000 cells/well for immunocytochemistry and 150,000 cells/well for luciferase assay), 12-well dishes (250.000 cells/well for molecular assays), 6-well dishes (500.000 cells/well for biochemical and molecular assays) or 60 mm diameter dishes (1.5-2×10^6^ cells/dish for subcellular fractionation and ChIP assays). Neurons were maintained in a humidified incubator at 37°C and 5% CO2.

### ChIP sequencing and gene expression analyses

Cortical and hippocampal neurons (20×10^6^ per ChIP condition) from mouse embryos (C57BL/6; E15.5; 11 DIV) were treated with vehicle (DMSO) or KCl (55 mM) plus forskolin (FSK, 20 μM). Chromatin was crosslinked, fragmented by sonication and immunoprecipitated using a rabbit monoclonal anti-CRTC1 antibody (#2587S, Cell Signaling, Danvers,USA), anti-CREB (#9197, Cell Signaling) and rabbit polyclonal anti-CRTC2 (#12497-1-AP, Proteintech), as described (Parra-Damas et al., 2017a; Parra-Damas et al., 2017b). Specific immunoprecipitation of endogenous CREB, CRTC1 and CRTC2 using these antibodies was verified by biochemical analyses (data not shown). For CRTC1/2 ChIP-Seq experiments, frozen PFA cross-linked cell pellets were gently resuspended and incubated in 1.5 mM ethylene glycol-bis(succinic acid N-hydroxysuccinimide ester; EGS) in PBS for 15 min and washed twice in PBS, to further cross-link CRTCs to DNA before chromatin shearing. Two independent ChIP experiments per condition were used for library preparation and sequencing. DNA libraries were prepared from 2.5 ng of immunoprecipitated or input DNA using the NEBNext Ultra DNA Library Prep Kit for Illumina (#E7370, New England Biolabs, Ipswich, USA) and sequenced (40M bp sequencing depth, 1 x 50 bp) on an Illumina HiSeq2500 instrument, following standard ChIP-seq guidelines (Kharchenko et al., 2008). ChIP-seq data analysis, including alignment to the reference genome, background (Input) filtering and peak calling, was done using the “mosaics” and “ChIPseeker” packages in Bioconductor, following established ChIP-seq guidelines (Kharchenko et al., 2008; Yu et al., 2015). PCA and differential binding analysis of the identified peaks was performed using the “DiffBind” package in Bioconductor. DNA libraries and ChIP-seq data were validated by quantitative real time RT-PCR (qRT-PCR) using specific primers (**Table S3**) (Parra-Damas et al., 2017b). **ChIP-seq data from this study are deposited in the Gene Expression Omnibus (GEO) database (accession number: GSE131395).**

### Gene expression analyses

Quantitative real-time RT-PCR was performed as described (Parra-Damas et al., 2017b). RNA from cultured hippocampal neurons or hippocampal tissue was purified with the PureLink RNA Mini Kit (Thermo Fisher Scientific). RNA integrity number (RIN) was measured using the Agilent 2100 bioanalyser (Agilent Technologies). RNA (1 μg; RIN > 8.0) was reverse-transcribed in 50 μl of a reaction mix containing 1 μM of Oligo (dT) primers, 1 μM random hexamers, 0.5 mM dNTP, 0.45 mM DTT, RNAseOut (10 units) and SuperScript^TM^ II reverse transcriptase (Thermo Fisher Scientific) at 25°C for 10 min, 42°C for 60 min and 72°C for 10 min. Amplification was performed by duplicates in 3-5 samples using specific primers (**Table S3**) in an Applied Biosystems 7500 Fast Real-Time PCR system (Thermo Fisher Scientific). Data analysis was performed by the comparative ΔCt method using the Ct values and the average value of PCR efficiencies obtained from LinRegPCR software. Gene expression was normalized to *Gapdh* for cultured neurons or the geometric mean of *Gapdh*, hypoxanthine guanine phosphoribosyl transferase (*Hprt*) and peptidylprolyl isomerase A (*Ppia*) for brain samples (Parra-Damas et al., 2017b).

### In vivo adeno-associated viral injections

Adeno-associated virus (AAVrh.2/10) containing *GFP* or *Crtc1-myc* under the cytomegalovirus (CMV) promoter (AAV-GFP and AAV-CRTC1-IRES-GFP) were generated as described (Parra-Damas et al., 2014). For *in vivo Crtc1* silencing, CRTC1 Short hairpin (Sh) RNAs known to target murine *Crtc1* (España et al., 2010), were synthesized and subcloned into AAV2/10 vector downstream of the histone 1 (H1) promoter followed GFP gene under the rous sarcoma virus (RSV) promoter. AAV-GFP, AAV-*Crtc1-myc*, AAV-Sh Scramble and AAV-Sh*CRTC1* (3 μl; 5.1×10^11^gc/ml; 0.5 μl/min) were injected bilaterally into the dorsal hippocampus of isofluorane-anesthetized mice. The sterotaxic injection coordinates were (in mm) as follows: anteroposterior: -2.0 from Bregma; mediolateral: ±1.8 from Bregma; ventral: -1.8 from dural surface, according to Paxinos and Franklin’s brain atlas. We used 4.5-5 month-old male mice (C57BL/6 background; n=6-8 mice/group) at the time of injection, and mice were analyzed and sacrificed 1.5 months after injection.

### Biochemical analysis

Cultured hippocampal neurons from mouse embryos (E15.5; C57BL/6) were non-transduced or transduced with scramble, CRTC1-myc or *Crtc1* ShRNA lentivirus (2 ip/cell). For analysis of GluN1 phosphorylation, cortical neurons (19-20 DIV) were incubated with tetrodotoxin (1 μM; Tocris), and then treated with vehicle or GF-109203X hydrochloride (10 μM; Sigma) before adding phorbol 12-myristate 13-acetate (PMA; 5 μM; Sigma) or bicuculline (50 μM; Sigma) and 4-AP (2.5 mM; Sigma) for 15 min. For transcription inhibition experiments, cortical neurons (20 DIV) were incubated with Actinomycin D (ActD; 2 μg/ml; Thermo Fisher) for 1 h and then treated with Bic/4AP for 15 min/1 h. Mouse neurons or hippocampi were lysed in cold-lysis buffer (50 mM Tris hydrochloride, pH 7.4, 150 mM NaCl, 2.5 mM ethylenediaminetetraacetic acid [EDTA], 1 mM Na_3_VO_4_, 25 mM NaF, 0.1% sodium dodecyl sulfate [SDS], 1% NP-40, 0.5 % Na-deoxycholate) containing protease and phosphatase inhibitors (Roche España). Proteins were quantified with the BCA protein assay kit (Thermo Fisher Scientific), resolved on SDS-polyacrylamide gel electrophoresis, transferred to PVDF membranes and blotted with the antibodies: anti-CRTC1 (1:10,000) and pGluN1 (Ser890, Ser897) (1:1,000; Cell Signaling); c-myc (1:1,000; Santa Cruz Biotechnology); GluA1 (1:2,000), GluN2B (1:2,000), PSD95 (1:2,000), pGluA1 (Ser845), GluA2, GluN1 and GluN2A (1:1,000; Merck-Millipore); synaptophysin (1:100,000) and β-tubulin (1:20,000; Sigma-Aldrich); and GAPDH (1:100,000; Thermo Fisher Scientific). Protein bands were imaged and quantified with a ChemiDoc MP System and Image Lab Software 5.2.1 (Bio-Rad) and normalized with GAPDH or β-tubulin.

### Cell surface biotinylation and subcellular fractionation

For biotinylation assays, hippocampal neurons (postnatal day 0) were incubated in cold PBS supplemented with 1 mM CaCl_2_ and 0.1 mM MgCl_2_ pH 7.4 (PBS-Ca-Mg) containing 1mg/ml EZ-link-sulfo-NHS-LC-biotin (Thermo Fisher) for 30min at 4°C. Neurons were rinsed in PBS-Ca-Mg plus 0.1 M glycine and lysed in 1% Triton X-100 and (in mM): 50 HEPES pH 7.5, 50 NaCl, 10 EDTA, 10 EGTA, 1 NaVO_3_, 50 NaF, 25 NaPPi, 1 β-glycerophosphate, 1 PMSF containing protease and phosphate inhibitors. After centrifugation (10,000 x *g,* 20 min), biotinylated proteins were isolated overnight at 4°C with Neutravidin Ultralink Resin agarose beads (Thermo Fisher). Proteins were eluted with 2X SDS-PAGE buffer (100 °C, 5 min) and loaded onto SDS-PAGE gels.

For subcellular fractionation, hippocampal neurons were lysed in HEPES-sucrose buffer containing (in mM): 4 HEPES pH 7.4, 320 sucrose, 1 NaVO_3_, 1 NaF, 0,1 NaPPi, 0,1 glycerophophate supplemented with protease and phosphatase inhibitors, and spun (1,000 x *g,* 10 min). Supernatant was spun twice for 15 min at 10,000 x *g*. The pellet containing the synaptosomal plasma membrane was resuspended in HEPES buffer and centrifuged (25,000 x *g,* 20 min). The pellet was resuspensed in HEPES-sucrose buffer, loaded in a discontinuous sucrose gradient and centrifuged (150,000 x *g,* 2 h). The pellet was resuspended in HEPES-EDTA buffer containing 0.5% Triton X-100, 50 mM HEPES, 2 mM EDTA and protease and phosphatase inhibitors, and centrifuged (32,000 x *g,* 20 min) to obtain the postsynaptic density fraction. For hippocampus subcellular purification, postsynaptic fractions were isolated and purified by sucrose gradient fractionation. Analysis of proteins was determined using Western blotting as above.

### Immunohistochemical and immunofluorescence staining

Mice were anesthetized with isofluorane and perfused transcardially with PBS and 4% phosphate-buffered paraformaldehyde before paraffin embedding. Sagittal brain sections (5 μm) were heated for 10 min in citrate buffer and incubated with rabbit antibodies: c-myc (sc789; 1:500; Santa Cruz Biotechnology), anti-CRTC1 (1:300; Cell Signaling) followed by AlexaFluor-488/594-conjugated goat IgGs (1:400) and Hoechst (1:10,000; Thermo Fisher Scientific). Images (20x; zoom 0.5 or 40x; zoom 0.5) were captured with a Carl Zeiss Axio Examiner D1 LSM700 laser scanning microscope and analyzed with ImageJ. For immunocytochemistry assays, non-infected or lentiviral transduced hippocampal neurons (20 DIV) were pretreated with tetrodotoxin (16 h) and then incubated with vehicle, PMA or bicuculline/4-AP for 1 h. Neurons were fixed and incubated overnight with mouse anti-GluN1 (1:200; Sigma), rabbit anti-PSD95 (1:200; Cell Signaling) and/or chicken anti-GFP (1:1000; Abcam), and detected with AlexaFluor-647, 568 or 488 conjugated goat antibodies (1:300; Life Technologies), respectively.

### Golgi staining, diolistic labeling and image acquisition

Mice were perfused transcardially with PBS and 4% phosphate-buffered paraformaldehyde. The right hemisphere was processed for Golgi staining using the FD RapidGolgiStain TM Kit (FD NeuroTechnologies Inc.) and sectioned coronally (100 μm) as described (Enriquez-Barreto et al., 2014). CA1 segments of basal dendrites from the dorsal hippocampus, 45 μm away from the soma, were imaged every 0.5 μm using a Nikon Eclipse TE-2000E microscope (Enriquez-Barreto et al., 2014). Images were deconvoluted using Huygens Essential software adapted for bright-field images (Scientific Volume Imaging). For diolistic (DiI) labeling, we used the Helios Gene Gun System (Bio-Rad, Hercules) using a suspension containing 3 mg of DiI and 50 mg of tungsten particles (1.7 μm) (Thermo Fisher Scientific). Particles were delivered into 150 μm vibratome coronal sections at 80 psi. DiI-labeled basal dendrites of CA1 pyramidal neurons were imaged using a Leica SP5 confocal microscope (63x objective, zoom 5). For detection of DiI/CRTC1-myc-expressing neurons, sections were incubated with PBS/0.05% Triton X-100 and incubated for 48 h with rabbit anti c-myc antibody followed with Alexa 488-conjugated secondary antibody and Hoechst (1:300; Thermo Fisher Scientific). DiI/GFP- and DiI/c-myc-positive neurons were used for basal dendrites segment’s image acquisition and analysis as described (Enriquez-Barreto et al., 2014).

### Electrophysiological recordings

Electrophysiological recordings were performed in 6 month-old male mice (C57BL/6; n= 4-5 per group) intrahippocampal injected at 4.5-5 months with AAVs and tested for contextual fear memory. Hippocampal slices (350 mm) were obtained under ice-cold solution I consisting of (in mM): 124 NaCl, 2.69 KCl, 1.25 KH_2_PO_4_, 2 MgSO_4_, 1.8 CaCl_2_, 26 NaHCO_3_, and 10 glucose (pH 7.2, 300mOsm) (Andrade-Talavera et al., 2016). Slices were continuously oxygenated, perfused and maintained at room temperature (22–25°C). Field excitatory postsynaptic potentials (fEPSPs) recorded in the CA1 region were evoked at 0.2 Hz by a monopolar stimulation electrode placed in the “stratum radiatum” using brief current pulses (200 μs, 0.1–0.2 mA). Long-term potentiation was induced by a TBS protocol consisting in 5 episodes of 10 train stimulus at 5 Hz, each one with 4 pulses at 100 Hz. Recordings lasted 180 min after LTP induction. Long-term depression was induced by a 1 Hz stimulus protocol during 30 min. In paired-pulse experiments, two consecutive stimuli separated by 40 ms were applied. Data were filtered at 3 kHz and acquired at 10 kHz. For basal synaptic transmission, a stimulus-response curve (40-160 μA, mean of 12 fEPSPs) was compiled for each mouse. For the recording of AMPA and NMDA receptor-mediated currents, CA1 hippocampal neurons were recorded with a patch-clamp amplifier (Multiclamp 700B), and data were acquired using pCLAMP 10.2 software (Molecular Devices). Patch electrodes were pulled from borosilicate glass tubing, and had resistances of 4–7 MΩ when filled with (in mM): caesium chloride, 120; HEPES, 10; NaCl, 8; MgCl_2_, 1; CaCl_2,_ 0.2; EGTA, 2 and QX-314, 20 (pH 7.2–7.3, 290 mOsm L^−1^). Cells were excluded from analysis if the series resistance changed by more than 15% during the recording. NMDA currents were recorded at +40 mV in the presence of bicuculline (20 μM) and NBQX (10 μM), and AMPA currents were recorded at -65 mV in the presence of D-AP5 (50 μM) and bicuculline at the same position and stimulating intensity across different cells. At the same stimulating intensity, no high variability in current amplitudes was observed as determined by the I/O curves. Ten to twenty traces were averaged at each holding potential. The AMPA/NMDA ratio was calculated by dividing the peak of AMPA receptor current at -65 mV by the amplitude of NMDA receptor current measured at 50 ms at a holding potential of +40 mV.

### Image morphological analyses

Spine density was measured manually using ImageJ Plugin Cell Counter (Enriquez-Barreto et al., 2014). For morphological analysis, spines with or without (stubby spines) a clear neck were classified, and head area was manually measured using ImageJ software. Briefly, a single image from the stack was selected for analysis and the scale was set. Each spine was selected, duplicated, rescaled, and thresholded. The “Polygon selections” tool was used to define the perimeter of the head, splitting it from the neck or the dendritic shaft. A distribution analysis of the head area of necked spines was performed and data were fit to a normal Gaussian distribution to set the mean value as a reference. Spines were divided in small (head areas < mean value), large (head areas > mean value) or non-neck stubby spine categories (Enriquez-Barreto et al., 2014). The number of analyzed animals, dendrites and spines are described in **Tables S1 and S2**.

For immunofluorescence imaging analysis, images were captured with a Leica SP5 confocal microscope (63x objective, zoom 3) and analyzed using Imaris 8 software (Bitplane AG, Zurich, Switzerland). GluN1 (> 0.4 μm diameter) and PSD95 (> 0.7 μm diameter) positive puncta were considered for quantitative analyses in multiple primary and secondary dendrites.

### Behavioral studies

For contextual fear conditioning, 6-month-old male mice were placed into the conditioning chamber (15.9 x 14 x 12.7 cm; Med Associates) for 3 min and foot-shocked (1sec/0.8 mA). Fear memory was measured as freezing behavior, considered as complete absence of movement except respiration, before (context), immediately (immediate freezing) or 24 h after the footshock using Video Freeze Software (Med Associates, Fairfax, USA) (Parra-Damas et al., 2017a).

### Statistical analysis

Statistical analysis was performed using Student’s t test or one- or two-way analysis of variance (ANOVA) for multiple comparisons followed by Bonferroni, Tukey’s or Sidak’s *post hoc* tests. Behavioral results were compared using two-way ANOVA and Scheffe’s S *post hoc*. The Kolmogorov-Smirnov test was used to compare dendritic spine areas and neck lengths. Electrophysiological results were analyzed using the Clampfit 10.2 software (Molecular Devices, Sunnyvale, USA) and statistical comparisons were performed using Student’s t-test. Significant levels were considered as follows-unless than indicated-: \**P* < 0.05, \*\**P* < 0.01, and and \*\*\**P* < 0.001 or 0.0001. All statistics are included in the text and/or figure legends. Data are represented as mean ± SEM and analyzed using Prism 6.0 software (GraphPad, San Diego, USA).

